# The effects of vertical transmission on a spatially-structured host-parasite model

**DOI:** 10.64898/2026.01.22.701030

**Authors:** Jack Woodruff, Alex Best

**Affiliations:** School of Mathematical and Physical Sciences, University of Sheffield, Sheffield, UK

**Keywords:** Vertical transmission, Spatial structure, Pair approximation, Horizontal transmission, Trade-off

## Abstract

Vertical transmission of an infectious disease from parent to offspring is a common transmission route in many systems. Here we investigate the dynamics of a pathogen with both horizontal and vertical transmission within a spatially structured population. We introduce a lattice model with a pair approximation that includes both local and global transmission and reproduction. We find that vertical transmission can determine pathogen invasion and reduce the horizontal transmission rate required for invasion. When the majority of transmission and reproduction is local, vertical transmission can destabilise a host population to cause limit cycles. Given the advantages of a pathogen having both horizontal and vertical transmission routes, we extend the model to investigate the likelihood a mutant strain with both transmission modes will outcompete a resident strain with only horizontal transmission. When there is no trade-off the mutant always invades and when there is a trade-off with horizontal transmission, the mutant emerges when the cost to the horizontal transmission rate is not too large. Depending on how the mutant appears within the host population, it may have an initial advantage over the resident strain even if it cannot outcompete in the long-term. Our work demonstrates the potential importance of vertical transmission within host-pathogen dynamics.

## 1 Introduction

Vertical transmission occurs when a pathogen is transmitted from parent to offspring at or prior to birth (Burrell et al. 2017; Fine 1975). Vertical transmission can be termed perfect if all offspring of an infected host are born infected, and imperfect if only a proportion of the offspring are born infected. For plant pathogens a vertically-transmitted disease is spread via seeds, and can cause outbreaks of disease (Dombrovsky and Smith 2017; Maule and Wang 1996; Pagán 2022). For animals, depending on the reproductive processes, the virus may be passed to offspring during fetal development, egg development, birth or postnatally, and the virus can come from either parent (Mims 1981). Notable human diseases that are vertically transmitted include HIV/AIDS (Cardenas et al. 2023) and Hepatitis B (Riches et al. 2025). The other main transmission mode is horizontal transmission, when an infected host passes disease to a susceptible host already in the population, such as via aerosol, water or sexual contact (Antonovics et al. 2017; Fine 1975). It is common for a pathogen to have a mixture of both transmission modes, even if transmission via one mode is relatively rare compared to the other (Ebert 2013). Theory shows that vertical transmission allows pathogens to persist in a wider range of ecological conditions (Altizer and Augustine 1997).

Mathematical models of vertical transmission are somewhat under-represented compared to horizontal transmission models. Where they do exist, analysis of mean-field models with vertical transmission have given insights into the influence of this transmission mode on population dynamics (Altizer and Augustine 1997; Busenberg et al. 1983; Gao and Hethcote 1992; Lipsitch et al. 1995), as well as provided insights into specific diseases including Dengue (Adams and Boots 2010), HIV/AIDS (Naresh et al. 2006) and Hepatitis B (Anley et al. 2023). If a pathogen reduces the net growth of the host, it cannot invade via vertical transmission alone; horizontal transmission is required (Busenberg and Cooke 1993; Lipsitch et al. 1995). In contrast, a pathogen with only horizontal transmission is able to invade (Kermack and McKendrick 1927), but when vertical transmission is present it can be a determinant in pathogen invasion (Busenberg et al. 1983). When both modes are present, the two transmission routes can have different roles in the course of a disease outbreak. While horizontal transmission may be critical during the early stages of an outbreak, vertical transmission can have a more significant role once the disease is established (Lipsitch et al. 1995). Vertical transmission can also allow the pathogen to persist at lower levels of host density (Anderson and May 1981) and a wider range of ecological conditions compared to if the pathogen had only horizontal transmission routes (Altizer and Augustine 1997). In the case of perfect vertical transmission and no recovery, an outbreak could lead to total infection of a population (Lipsitch et al. 1995). For pathogens with both horizontal and vertical transmission routes, theoretical and experimental studies have considered the possibility of a trade-off between the two transmission modes (García-Ordóñez and Pagán 2024; Magalon et al. 2010; Stewart et al. 2005; Turner et al. 1998; Zilio et al. 2018). In a mean-field model with such a trade-off, bistability has been found, whereby depending on the environment, a pathogen can evolve high horizontal transmission and virulence with low vertical transmission or low horizontal transmission and virulence with high vertical transmission (van den Bosch et al. 2010).

The majority of ecological interactions are subject to spatial structure, particularly the transmission of disease, which often occurs through direct contact between susceptible and infected hosts. Spatial structure is particularly important in systems where individual hosts are static, such as plants. The importance of including spatial structure within a model is demonstrated by the fact that spatially structured models behave differently to mean-field counterparts. For example, limit cycles and disease-driven extinction can occur (Best et al. 2012; Webb et al. 2007a,b). Spatial structure can be introduced into models through a variety of methods, one of which is the lattice approach with a pair approximation (Filipe and Gibson 1998; Matsuda et al. 1992; Satō et al. 1994). Using this approach, recent research has investigated how spatial structure affects disease dynamics with only horizontal transmission by varying the proportion of local and global reproduction and transmission (Maltz and Fabricius 2016; Webb et al. 2007a; Wren and Best 2021). These studies have found that the proportion of local horizontal transmission has a larger impact than the proportion of local reproduction on the invasion boundary (Webb et al. 2007a). In contrast, the proportion of local reproduction has a larger affect on the occurrence of endemic limit cycles (Webb et al. 2007a).

While most theoretical studies with spatial structure assume only horizontal transmission, a few have included vertical transmission (Schinazi 2000; Silva et al. 2017; Su et al. 2019). In one such study of a lattice-based infectious disease model with perfect vertical transmission and no recovery, four steady states were identified (extinction, disease-free, endemic and total infection) and limit cycles were not found (Silva et al. 2017). Given that the birth rate is sufficient enough that extinction does not occur, the fecundity of infected hosts and the horizontal transmission rate are key parameters in determining disease invasion and whether total infection occurs (Su et al. 2019). For a stochastic model on a one-dimensional lattice where vertical transmission is imperfect and there is no recovery, it has been shown that if vertical transmission is below a threshold, then even for high horizontal transmission the disease will not persist (Schinazi 2000). When there is habitat loss, vertical transmission can play a more important role in whether the pathogen invades, due to the potential unavailability of susceptible hosts (Su et al. 2019). Despite these studies, a lattice-based deterministic model that allows for imperfect vertical transmission has not yet been considered, and it is unknown how imperfect vertical transmission will affect the behaviour of disease dynamics in spatially-structured populations.

Here we introduce such a model that also allows for both local and global horizontal transmission and reproduction, and therefore local and global vertical transmission. In Section 2, the model is introduced. Section 3 presents the key results from our analysis, including an extension of the model that allows for the emergence of a mutant with vertical transmission in a population where it is otherwise absent. Section 4 discusses the conclusions.

## 2 Model

### 2.1 Mean-Field Model

Consider a population comprised of susceptible (*S*) and infected (*I*) hosts, with a total population size *N* . For clarity, initially we assume there are no local interactions. The model is given by the following ordinary differential equations where dots represent the derivative with respect to time,

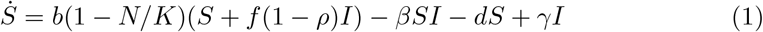

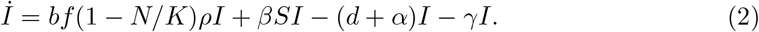

We assume horizontal transmission is density-dependant with co-efficient *β* and recovery rate *γ*. Reproduction is limited by carrying capacity *K* and occurs at rate *b* for susceptibles and *bf* for infecteds where *f* ∈ [0, 1] is fecundity. The probability of vertical transmission occurring is *ρ*. Natural death occurs at rate *d* with virulence, *α*, for infected hosts.

### 2.2 Pair-Approximation Model

We now introduce the spatial structure. Consider a square lattice, with periodic boundary conditions, where each site is empty (0) or occupied by a susceptible (*S*) or infected (*I*) host (Matsuda et al. 1992; Satō et al. 1994). Interactions are local if they occur between a site and one of its four nearest-neighbours, where as interactions are global if they occur between two random sites. Local reproduction occurs when susceptible and infected hosts reproduce into neighbouring empty sites, this occurs at rates *bL*_*R*_ and *bf L*_*R*_ respectively, where 0 ≤ *L*_*R*_ ≤ 1, and hosts reproduce into random empty sites at rates *b*(1 − *L*_*R*_) and *bf* (1 − *L*_*R*_) respectively. Vertical transmission occurs locally at rate *ρbf L*_*R*_ and globally at rate *ρbf* (1 − *L*_*R*_), otherwise infected offspring are born susceptible. As in Eq. 1-2, we assume horizontal transmission is density dependant with it occurring locally between infected and susceptible neighbours at rate *βL*_*T*_ and globally at rate *β*(1 − *L*_*T*_) for 0 ≤ *L*_*T*_ ≤ 1. All other parameters are the same as for Eq. 1-2.

The probability that a site is in state *i* is defined as *P*_*i*_ for *i* ∈ {0, *S, I*}. Similarly, the probability that a neighbouring pair of sites are in states *ij* is defined as *P*_*ij*_ where *i, j* ∈ {0, *S, I*} . For each pair of sites in state *ij, q*_*i/j*_ is the conditional probability that for a randomly chosen *j*-site, a randomly chosen neighbour is an *i*-site, with *q*_*i/j*_ = *P*_*ij*_*/P*_*j*_. Similarly, *q*_*k/ij*_ is the conditional probability that given an *ij*-pair, a randomly chosen neighbour of the *i*-site is a *k*-site.

Using these definitions, we define a system with both local and global reproduction and transmission, where transmission occurs either horizontally or vertically, using the following ordinary differential equations,

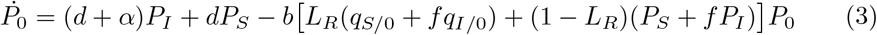

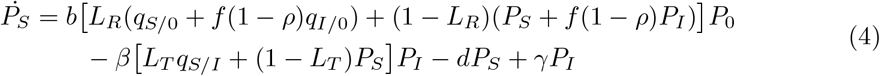

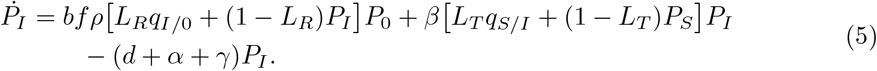

This system is not closed because the singlet equations depend upon the pair densities, due to the conditional probabilities. For example *q*_*S/I*_ = *P*_*SI*_*/P*_*I*_ . The dynamics of the pair probabilities are defined as,

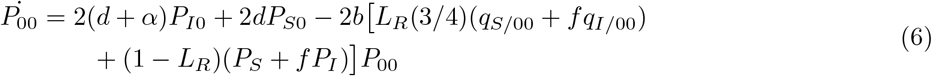

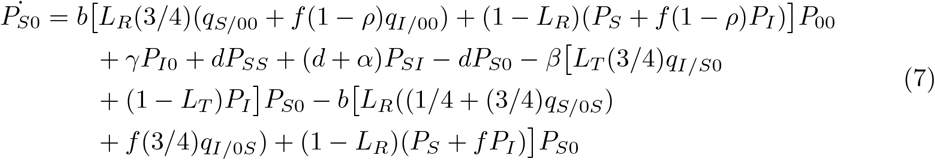

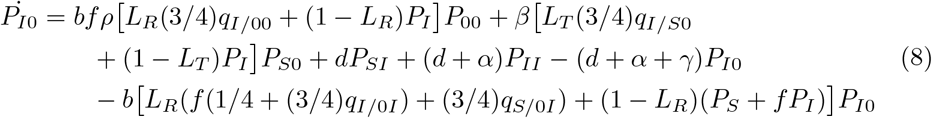

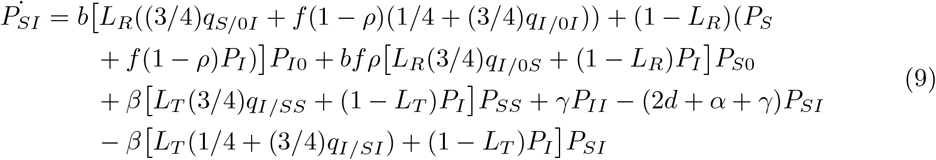

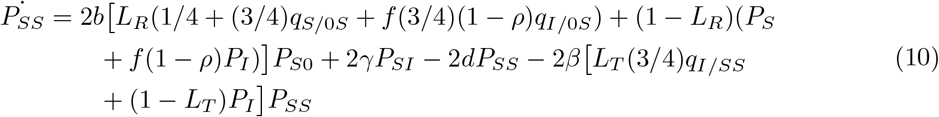

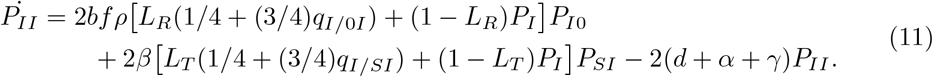

We also have that,

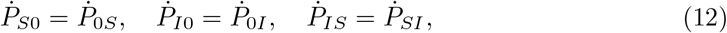

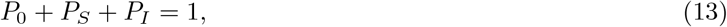

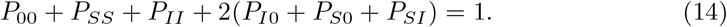

Within the conditional probabilities in Eq. 6-11 are the triplet densities. For example, *q*_*I/SI*_ = *P*_*ISI*_*/P*_*SI*_ . To close the system we use a pair approximation (Matsuda et al. 1992; Satō et al. 1994), such that,

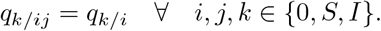

The validity of this approximation is shown by comparing the numerically simulated deterministic model to a spatially explicit stochastic simulation. The stochastic simulation is constructed using the Gillespie Algorithm (Gillespie 1977), with *τ* -leaping to increase the speed of each simulation run (Gillespie 2001).

## 3 Results

### 3.1 Invasion and Equilibria

For the fixed parameters in Fig. 1, there are three possible outcomes of the model: extinction, disease-free and endemic. Limit cycles do not occur for the parameter values used here, but are found for alternative parameter regions considered later in this section. Disease-driven extinction has been observed within spatial models with density dependent transmission. However, total castration or extremely low levels of infected reproduction are required for this steady states existence (Webb et al. 2007b). To observe the effects of vertical transmission, a sufficient level of infected reproduction is required, thereby eliminating the possibility of this steady state within our analysis.

**Fig. 1:**
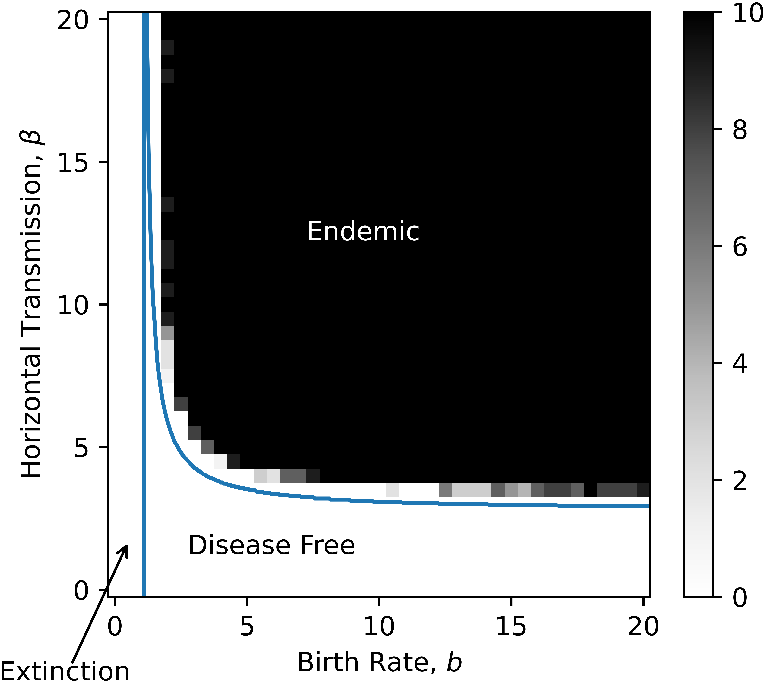
Equilibrium classification diagram in the *b*-*β* parameter space. The solid lines represent the invasion and extinction thresholds derived from numerical simulations of the deterministic model. The black and white shading shows the number of ten stochastic runs that resulted in pathogen invasion. The fixed parameters are *f* = 0.5, *d* = 1, *α* = 0.5, *γ* = 1.5, *ρ* = 0.75, *L*_*R*_ = *L*_*T*_ = 0.5. For the deterministic model the initial conditions are (*P*_0_, *P*_*S*_, *P*_*I*_, *P*_00_, *P*_*S*0_, *P*_*I*0_, *P*_*SI*_, *P*_*SS*_, *P*_*II*_) = (0.1, 0.8, 0.1, 0.01, 0.08, 0.01, 0.08, 0.64, 0.01). For the stochastic model, a 32x32 lattice began with 103 empty, 818 susceptible and 103 infected sites

The probability of vertical transmission can determine whether the pathogen invades. The minimum value of *ρ* for which this occurs is the critical probability of vertical transmission, *ρ*_*c*_. Focussing close to the invasion boundary, Fig. 2b demonstrates that in more localised systems, *ρ*_*c*_ increases, with *L*_*T*_ having a greater affect compared to *L*_*R*_. For this parameter set, invasion does not occur in the most localised system.

**Fig. 2:**
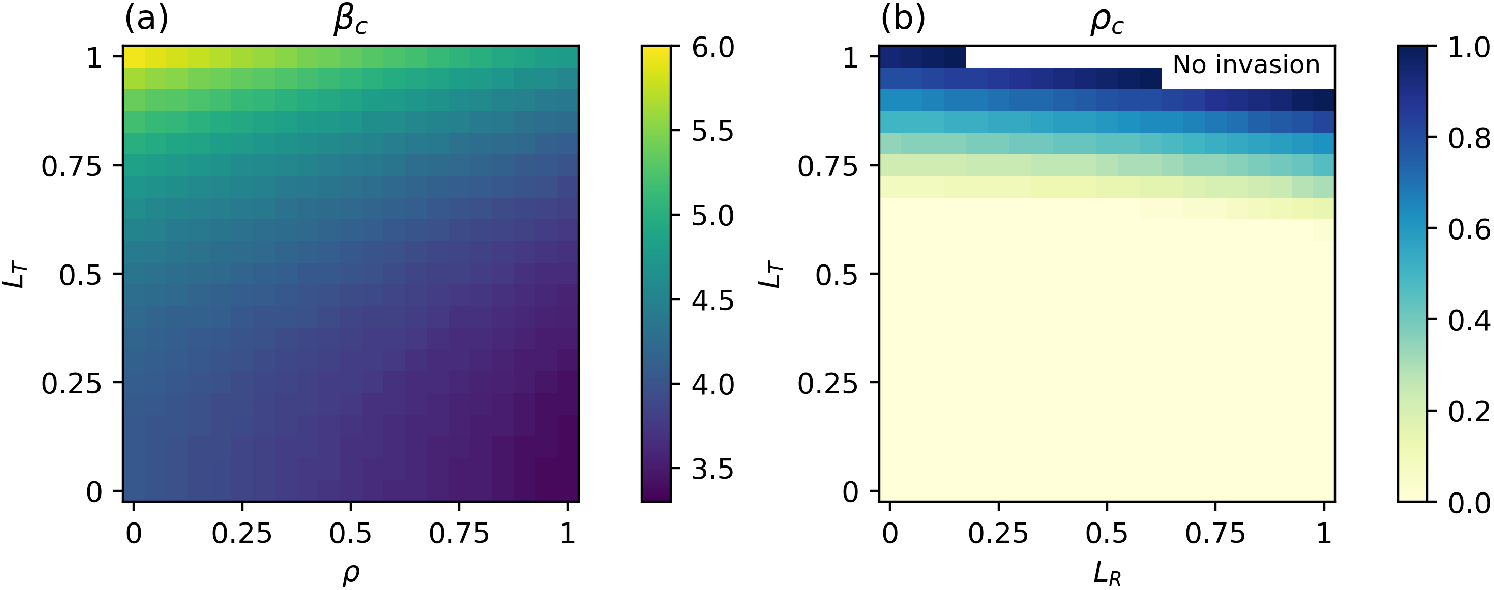
(a) Heatmap showing how *L*_*R*_ and *ρ* effect the critical horizontal transmission rate, *β*_*c*_. (b) Heatmap showing how *L*_*R*_ and *L*_*T*_ affect the critical probability of vertical transmission, *ρ*_*c*_. The fixed parameters are *b* = 4, *f* = 0.5, *d* = 1, *α* = 0.5, *γ* = 1.5 and (a) *L*_*R*_ = 0.5 (b) *β* = 4.6 with initial conditions as in Fig. 1

To compare the horizontal and vertical transmission routes, we look at the number of new cases over time, *C*_*h*_(*t*) and *C*_*v*_(*t*), for horizontal and vertical transmission respectively. Similarly to Lipsitch et al. (1995), these are defined as,

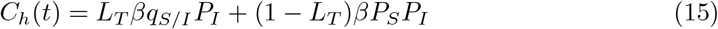

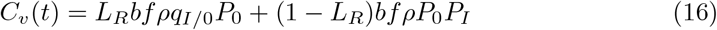

Previous work has shown that in mean-field models with vertical transmission, there are parameter values that result in only vertical transmission contributing to new infections at the steady state (Lipsitch et al. 1995). One such set is shown in Fig. 3a and 3c. However, in the equivalent spatial model this does not happen. As shown in Fig. 3d, horizontal transmission contributes new cases at the steady state. In the spatially structured model there are a higher number of susceptible hosts at the endemic fixed point, thus increasing *C*_*h*_ and lowering *C*_*v*_.

**Fig. 3:**
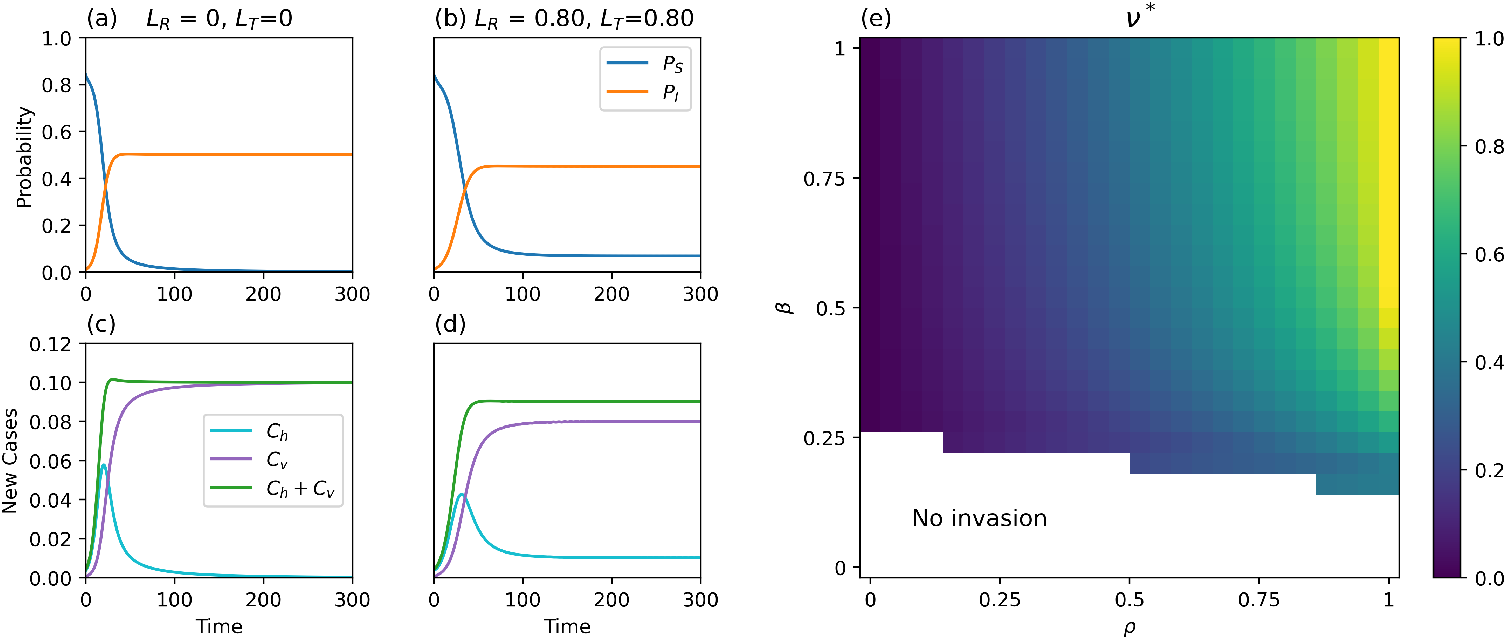
(a) and (b) show the time evolutions of the model with (c) and (d) showing the corresponding new cases over time respectively. (e) shows how the proportion of vertical transmission at the steady state, *ν*^∗^ varies in the *ρ*-*β* parameter space. The fixed parameters are taken from Lipsitch et al. (1995) with *f* = 2*/*3, *α* = 0.1, *γ* = 0, *ρ* = 1 with (a-d) *b* = 0.6 and *β* = 0.42. The spatial parameters are (a) *L*_*T*_ = *L*_*R*_ = 0 and (b,e) *L*_*R*_ = *L*_*T*_ = 0.8. For both cases the initial conditions are (*P*_0_, *P*_*S*_, *P*_*I*_, *P*_00_, *P*_*S*0_, *P*_*I*0_, *P*_*SI*_, *P*_*SS*_, *P*_*II*_) = (0.15, 0.84, 0.01, 0.005, 0.145, 0., 0.01, 0.685, 0)

Consider a time point, *t*_1_, that is sufficiently large so that the system has reached a steady state. The proportion of vertical transmission at the steady state is given by

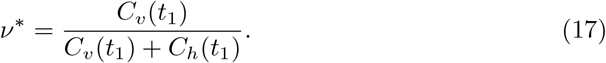

Fig. 3e, shows that, as expected, increasing *ρ* results in more vertical transmission at the steady state. Perhaps unexpectedly, increasing *β* also causes *ν*^∗^ to increase. Since increasing *β* reduces the susceptible population at the steady state, *C*_*h*_ is also lower at the steady state, and therefore *ν*^∗^ increases. Fig. 3d shows that initially *C*_*h*_ is higher than *C*_*v*_, suggesting that the invasion of the pathogen depends more on new cases arising via horizontal transmission. This explains why in Fig. 2b we see *L*_*T*_ having a larger affect on *ρ*_*c*_ compared to *L*_*R*_, even though *L*_*R*_ is more closely related to *ρ*.

As well as the steady states shown in Fig. 1, the model also gives rise to limit cycles. They only occur in certain parameter regions, when *f* is very small (see Appendix A). It is also required that *α* and *γ* are small, and *b* and *β* large. As with other parameters, the probability of vertical transmission can determine whether limit cycles occur. When cycles do occur, decreasing *ρ* decreases the amplitude of cycles, as shown in Fig. 4. This suggests that susceptible offspring from infected hosts can stabilise cycles.

**Fig. 4:**
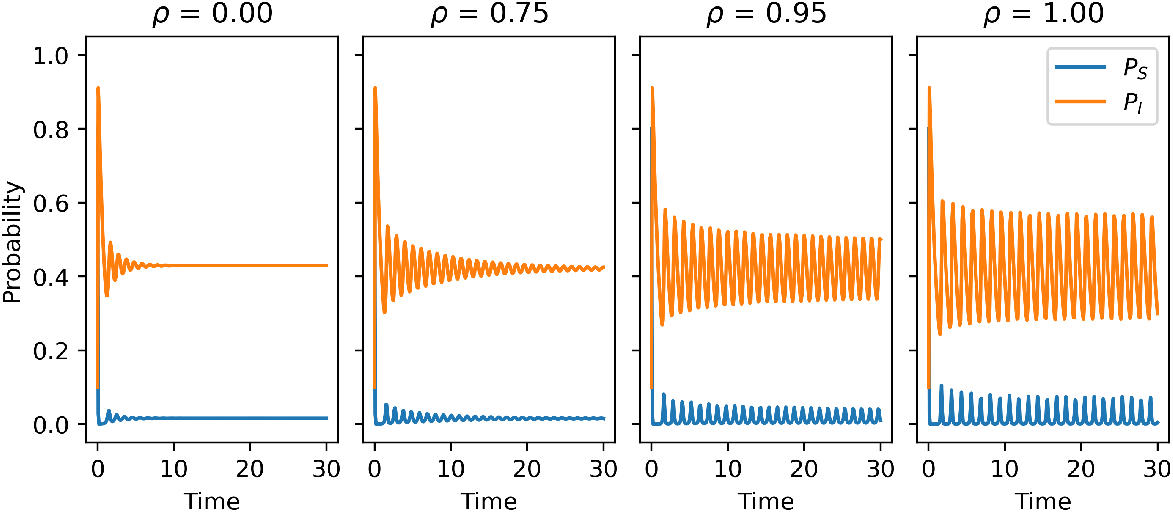
Time evolutions of the model for cases when dampened oscillations or limit cycles occur. The fixed parameters are *b* = 80, *f* = 0.001, *d* = 1, *α* = 0.05, *γ* = 0.0005, *β* = 80, *L*_*R*_ = *L*_*T*_ = 0.95 with *ρ* = {0, 0.75, 0.95, 1}. The initial conditions are the same as Fig. 1

Limit cycles occur when there are low levels of global transmission and reproduction, with a larger amplitude in more localised systems, as shown in Fig. 5. *L*_*R*_ has a larger influence on whether limit cycles occur compared to *L*_*T*_ . In Fig. 5, it is required that *L*_*R*_ ≥ 0.9. When reproduction is entirely local (*L*_*R*_ = 1), then limit cycles can continue to occur even when horizontal transmission is mainly global (*L*_*T*_ *<* 0.5).

**Fig. 5:**
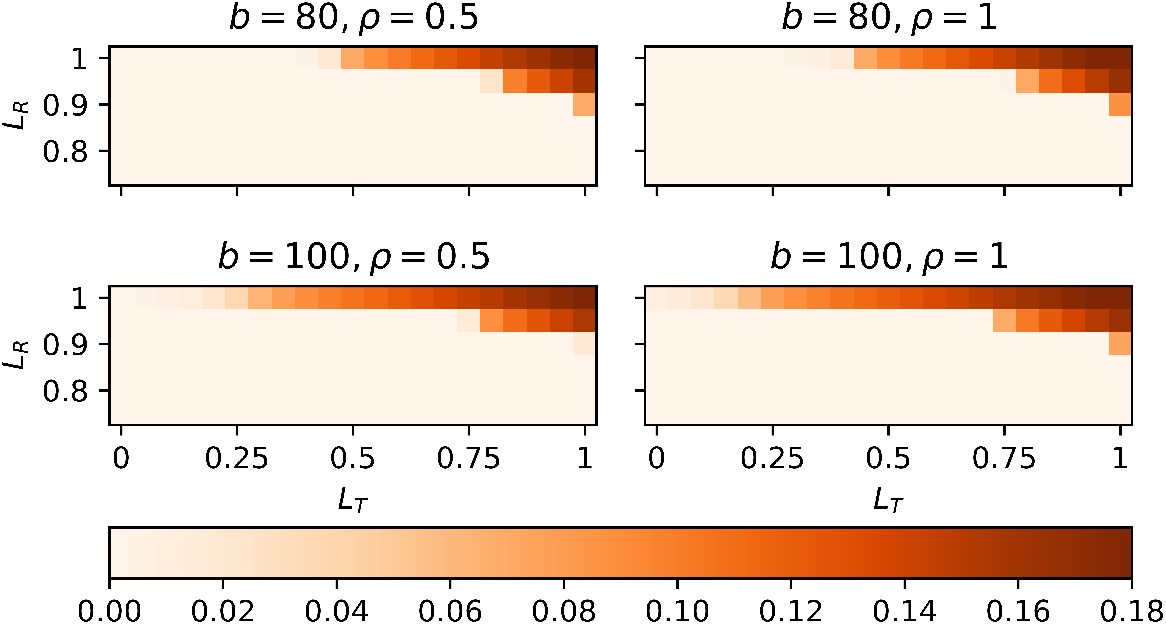
Heatmaps showing the amplitude of limit cycles. The fixed parameters are *f* = 1 *×* 10^−^5, *d* = 1, *α* = 0.05, *γ* = 0.0005 and *β* = 80. The initial conditions are the same as Fig. 1

However, once there is even a small amount of global reproduction, limit cycles only occur when the majority of transmission is local.

### 3.2 Invasion of a vertically transmitted strain

We now ask how easily a strategy of vertical transmission can emerge in a pathogen population that previously only had horizontal transmission. Consider a resident strain that is only transmitted horizontally, where *P*_*I*_ is the probability a site is occupied by a host infected with the resident strain. *P*_*V*_ is the probability a site is occupied by a host infected with the mutant strain that has vertical transmission as an additional transmission mode. *P*_*S*_ and *P*_0_ are as defined in Section 2. The pair probabilities are *P*_*ij*_ for *i, j* ∈ {0, *S, I, V*} . The system is defined by the following ordinary differential equations,

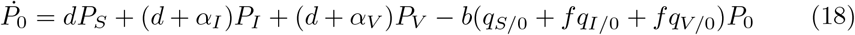

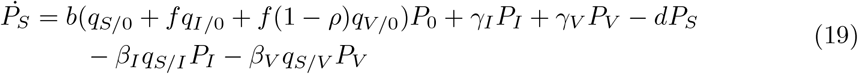

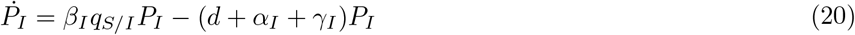

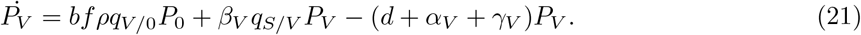

The parameters are as defined in Section 2, and, where applicable, the subscripts denote the strain corresponding to the parameter. The pair equations are in Appendix B defined by Eq. B4-B13. Once again, the system is closed using a pair approximation as outlined in Section 2 (Matsuda et al. 1992; Satō et al. 1994).

Initially, we assume the lattice is comprised of sites in states 0, *S* or *I*, and 10 that the system is at an endemic steady state expressed by 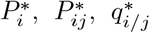 with 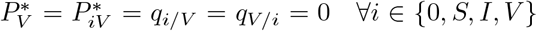 ∀*i* ∈ {0, *S, I, V*} . At time *t*_*M*_, a mutant is introduced into the population through one of three event types as defined in Table 1. Event type *S* is when during horizontal transmission the pathogen mutates. Event type *I* occurs when the pathogen mutates within an infected host. Biologically, we consider this type to be most likely. And for event type 0, a host with the mutant strain migrates into the population. We assume that the local environment of the mutant is the same as the site-type from which it mutated. Following the mutant event, the system is defined by 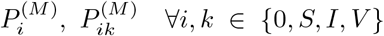 ∀*i, k* ∈ {0, *S, I, V*}. For event type *j*, the probability densities are expressed as,

**Table 1.** The three different event types for which a mutant may be introduced into the population. Note that by the pair approximation, we have that *q*_*i/V*_ = *q*_*i/V I*_ = *q*_*i/SI*_ = *q*_*i/S*_

| Event Type | Process | Local environment assumption $\forall i \in \{0, S, I, V\}$ |
| --- | --- | --- |
| S | $SI \rightarrow VI$ | $q_{i/VI} = q_{i/SI}$ |
| I | $I \rightarrow V$ | $q_{i/V} = q_{i/I}$ |
| 0 | $0 \rightarrow V$ | $q_{i/V} = q_{i/0}$ |

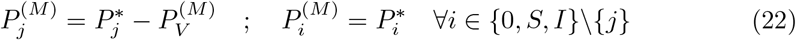

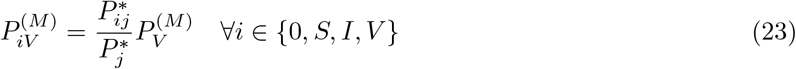

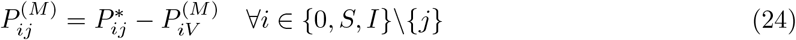

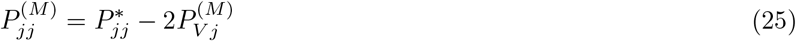

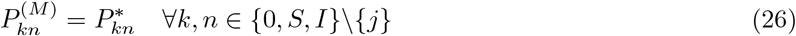

Following a mutation event, the mutant will either invade and replace the resident strain, invade and co-exist with the resident strain or die out. Initially, consider the case when *β*_*I*_ = *β*_*V*_, *α*_*V*_ = *α*_*I*_ and *γ*_*V*_ = *γ*_*I*_, such that there is no cost to vertical transmission. For all three event types, the mutant will always invade and either replace or co-exist with the resident strain. For event type *I*, if 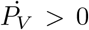 at *t* = *t*_*M*_ then the mutant will invade, which is always the case. For event types *S* and 0, the mutant will invade if 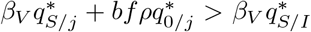 for event type *j* ∈ {0, *S*} . Through numerically simulating the time evolution of the model for 16525 parameter sets of which 9720 sets resulted in the initial resident strain invading, these inequalities were found to always hold for event types *S* and 0.

Given vertical transmission is expected to always emerge when cost-free, we now consider the potential for invasion when there is an associated cost. We assume that the mutant strain will have reduced horizontal transmission by assuming *β*_*V*_ = *ηβ*_*I*_ for *η* ∈ [0, 1). We also assume that virulence and recovery are the same for both strains (i.e. *α* = *α*_*I*_ = *α*_*V*_ and *γ* = *γ*_*I*_ = *γ*_*V*_).

For all three event types the long-term dynamics of the model are identical. Fig. 6a shows that for event type *I*, 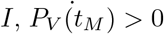 can be used to predict when the mutant will invade. However, for event types 0 and S, 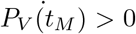 does not correctly predict the long-term dynamics of the model. Due to the mutant appearing in a different environment to other infected hosts, the mutant strain has a brief advantage causing initial growth, even if in the longer-term the mutant is unable to outcompete the resident strain. For example, in event type *S* the mutant is more likely to be surrounded by susceptibles compared to the event type *I* case. A similar logic applies to event type 0 as the mutant is more likely to have neighbouring empty-sites to reproduce into. A similar result was found in the case of a trade-off with virulence (see Appendix C). In both cases, there needs to be a fairly strong cost to vertical transmission for the mutant strain to not invade.

**Fig. 6:**
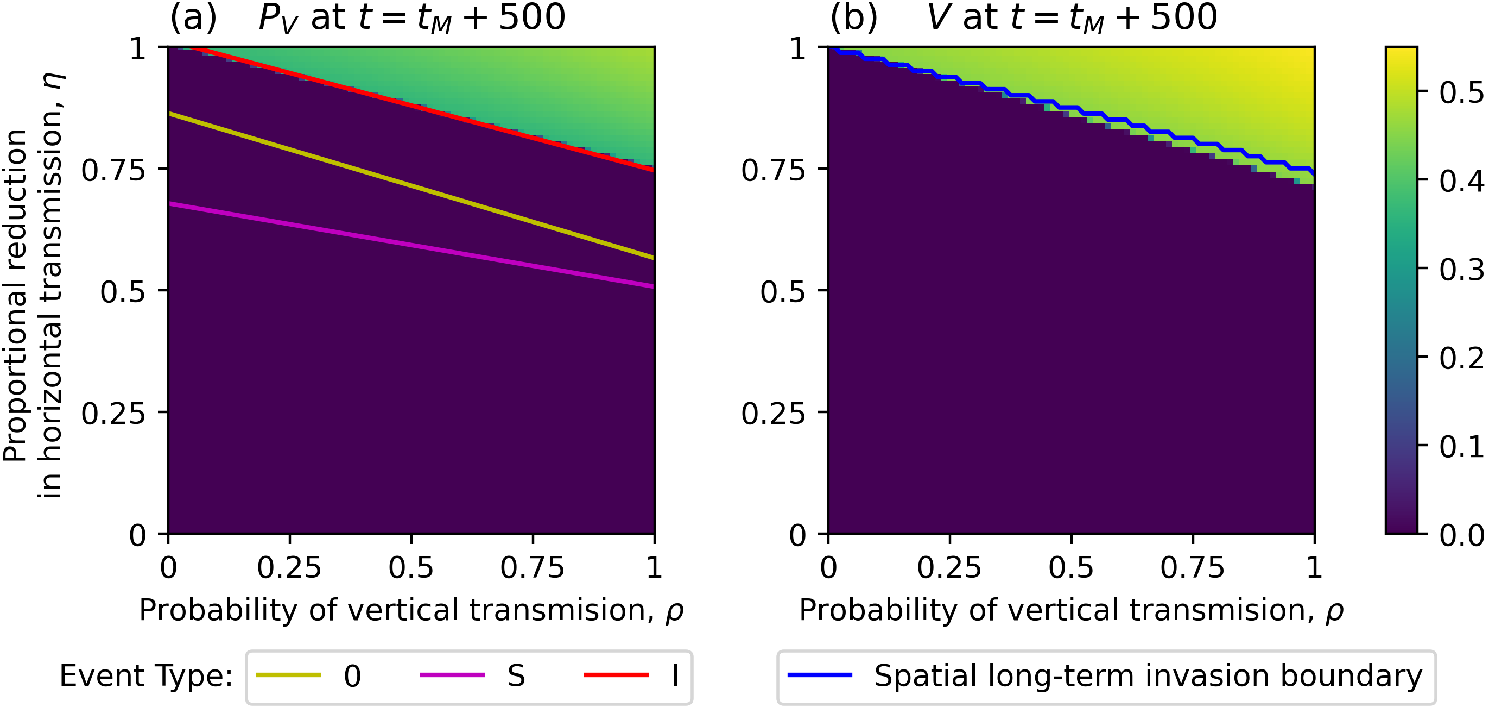
(a) Heatmap showing *P*_*V*_ at *t* = *t*_*M*_ + 500 for the spatial model. The solid lines represent the boundary where 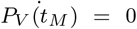. (b) Heatmap showing *V* at *t* = *t*_*M*_ + 500 for the mean-field model. The solid line represents the long-term invasion boundary in the spatial population. The fixed parameters for (a) and (b) are *b* = 10, *f* = 0.5, *d* = 1, *α*_*I*_ = *α*_*V*_ = 0.5, *γ*_*I*_ = *γ*_*V*_ = 1.5, *β*_*I*_ = 8, *β*_*V*_ = *ηβ*_*I*_, *t*_*M*_ = 500 and the additional parameter in (b) is *K* = 1. The initial conditions for (a) are (*P*_0_, *P*_*S*_, *P*_*I*_, *P*_*V*_, *P*_00_, *P*_*S*0_, *P*_*I*0_, *P*_*V* 0_, *P*_*SI*_, *P*_*SV*_, *P*_*IV*_, *P*_*SS*_, *P*_*II*_, *P*_*V V*_) = (0.1, 0.8, 0.1, 0, 0.013, 0.078875, 0.008125, 0, 0.082375, 0, 0, 0.63875, 0.0095, 0) with 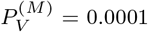 and for (b) are (*S, I, V*) = (0.8, 0.1, 0) with *V* (*t*_*M*_) = 0.0001

In the mean-field model, defined by Eq. B1-B3 in Appendix B.1, 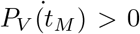 does accurately predict the long-term dynamics of the model. The mean-field model has a similar invasion threshold when compared to the spatial model, with a difference only occurring for higher *ρ*. Therefore, the mutant is less likely to invade in the spatial model. The final density of the mutant is lower in the spatial model.

## 4 Discussion

We have analysed a theoretical model of imperfect vertical transmission and its impact on host-parasite dynamics. One focus of our study has been the conditions that will allow for the initial emergence of vertical transmission. The full evolutionary dynamics of vertical transmission when traded off with horizontal transmission within a spatial model are currently unknown. Here, we have taken an initial investigation that includes the possibility of vertical transmission emerging as an additional transmission mode in a mutant strain. We found such a mutation to commonly be a favourable strategy, especially when there is no cost, in which case the mutant population will always grow, and either co-exist with or displace the resident strain. Biologically it is more likely that there is a cost to such a mutation, and this has been observed as a trade-off with either virulence (Cobos et al. 2019; Stewart et al. 2005) or horizontal transmission (Stewart et al. 2005; Turner et al. 1998; Zilio et al. 2018). We find that even if there is a cost, a mutant that can transmit vertically as well as horizontally is still likely to emerge. Notably, however, due to the spatial structure, the initial growth of the mutant strain does not always predict the long-term dynamics, depending on how the new strain appears. If it replaces a previously susceptible host - i.e. the mutation occurs upon infection - the mutant’s local environment is that of a susceptible host, and therefore different to that of the resident parasite strain. In spatially structured models, susceptible hosts that reproduce form clusters, so the newly emerged infectious mutant will have a higher availability of susceptible hosts compared to the resident strain. In some cases this advantage is short lived and the mutant is unable to persist in the long-term, due to its decreased horizontal transmission rate compared to the resident strain. Similarly, when a host infected with the mutant strain emerges in an empty site, perhaps via migration, initially it may have a larger number of empty sites to reproduce into compared to the resident parasite strain. This is an advantage because reproduction is a potential transmission event. When the pathogen mutates within an infected host, the mutant emerges with the same local environment as the resident strain so has no added advantage of having more susceptible hosts or empty sites available. In this case the initial growth of the mutant does accurately predict the long-term dynamics. Despite these early differences, for each mutation event the long term dynamics are identical, suggesting that the way the mutant emerges only influences the initial dynamics of the mutant strain. Compared with a mean-field population, the final density of the mutant strain is lower in the spatial population. This is to be expected because in spatial populations disease prevalence is lower when compared to the mean-field case (Webb et al. 2007a). The invasion boundaries of the spatial and mean-field populations differ only for higher levels of vertical transmission, suggesting spatial structure may only affect the initial dynamics when the mutation is for high vertical transmission.

We might therefore conclude that a mutant with vertical transmission as an additional transmission mode is likely to invade, which raises the question asked by Ebert (2013) as to why pathogens with multiple transmission modes are not observed more? Lab studies have shown that additional transmission routes are beneficial to pathogen invasion and persistence (Mangin et al. 1995). The lack of examples observed may be due to some unknown cost, trade-off or biological restriction not accounted for in current evolutionary studies. Negative trade-offs between vertical and horizontal transmission have been observed in both specific ecological contexts (Herre 1995; Jaenike 2000; Stewart et al. 2005; Turner et al. 1998; Zilio et al. 2018) and explored within mathematical models (Mangin et al. 1995; van den Bosch et al. 2010). However, there have been cases where the trade-off has been positive (Ebert 2013). Here we considered only the negative trade-off because in populations that are more spatially structured the trade-offs observed have been negative (Ebert 2013; Herre 1995; Jaenike 2000; Stewart et al. 2005).

Our results suggest that a mutant with any level of vertical transmission can persist in a spatially structured system, if the mutant strain has a horizontal transmission rate above a certain threshold. Therefore, a strain with low or high vertical transmission could invade. This does not rule out the possibility of the bistability observed in a mean-field model (van den Bosch et al. 2010) occurring within a spatially structured model. van den Bosch et al. (2010) found that a pathogen may evolve either high vertical transmission but low horizontal transmission and virulence, or low vertical transmission but high horizontal transmission and virulence. A natural extension of our analysis here would be to develop a full evolutionary model of vertical transmission in a spatially-structured population. This would allow a more direct comparison to mean-field evolutionary models that include vertical transmission within the tradeoff (Bernhauerová and Berec 2015; Lipsitch et al. 1996; van den Bosch et al. 2010).

As in the mean field model of the ecological dynamics with vertical transmission (Busenberg and Cooke 1993; Lipsitch et al. 1995), we find in the spatially structured model that horizontal transmission is required for pathogen invasion. While vertical transmission alone cannot result in pathogen invasion, when it is an additional transmission mode it does aid in invasion in both a mean-field (Busenberg et al. 1983) and spatial model. A pathogen with both vertical and horizontal transmission is able to persist in a wider range of ecological contexts, for example, when host density is lower (Ebert 2013; Lipsitch et al. 1995). This is reflected in our model where we observe that increasing the probability of vertical transmission reduces the minimum horizontal transmission rate required for invasion. Using the same methodology as Lipsitch et al. (1995), we demonstrated the different roles horizontal and vertical transmission play throughout an epidemic. In both mean-field and spatial contexts, horizontal transmission is dominant in the early stages and vertical transmission is dominant when disease prevalence is high. This is because as the susceptible host density lowers due to horizontal transmission, the number of infected hosts increases. A higher number of infected hosts results in a higher number of infections caused by vertical transmission. The swapping of dominant transmission modes between horizontal early on and vertical transmission at the steady state, suggests that different control strategies will be required depending on the epidemic stage (Aliabadi et al. 2011; Cardenas et al. 2023; Idigo et al. 2025; Veronese et al. 2021).

We also observe that increasing the probability of vertical transmission reduces the number of new cases caused by horizontal transmission in the later stages of an epidemic. When vertical transmission is absent, infected reproduction results in infected hosts gaining susceptible neighbours who can then be infected horizontally. However, when vertical transmission is present, infected reproduction may result in more infected neighbours which then reduces the availability of susceptible hosts. Higher horizontal transmission rates also result in a decreased availability of susceptible hosts, which in turn increases the proportion of vertical transmission at the steady state. Therefore, if disease prevalence is extremely high and susceptible host density is low, then vertical transmission is likely to be the primary driver behind new cases. In their mean-field model Lipsitch et al. (1995) present a case where only vertical transmission is contributing to new cases at the steady state due to there being no susceptible hosts remaining, and therefore total infection occurring. When local processes are included, the same parameter values result in a steady state where horizontal transmission is still contributing to new cases. Spatial structure is known to result in slower rates of disease spread (Keeling 1999), in turn leading to lower infected and higher susceptible densities compared to the mean-field (Webb et al. 2007a), leading to the potential for ongoing horizontal transmission.

Our model includes both local and global processes to allow for insight into how vertical transmission is affected by varying levels of spatial structure. Webb et al. (2007a) found that increasing the proportion of local transmission and reproduction results in a higher horizontal transmission rate required for invasion. We find that vertical transmission behaves similarly. While the level of local reproduction is closely tied to vertical transmission, the proportion of local horizontal transmission has a larger impact on the minimum probability of vertical transmission required for pathogen invasion. This is likely because local transmission is closely correlated to horizontal transmission, and in the early stages this is the dominant transmission mode so a key determinant in invasion.

As in previous studies of spatially-structure host-parasite dynamics (Webb et al. 2007a,b), we find that limit cycles occur when a high proportion of transmission and reproduction are local, and when infecteds’ fecundity is low. In particular, we observe that local reproduction has a larger effect on limit cycles compared to local horizontal transmission (Webb et al. 2007a). We have identified that vertical transmission can play a key role in the occurrence of limit cycles, where higher levels of vertical transmission are more likely to lead to cycles. Reducing the probability of vertical transmission results in more susceptible offspring from infected hosts, which Webb et al. (2007b) found to stabilise limit cycles. However, to observe the effects of vertical transmission, our model needs to include an adequate level of infected reproduction. Therefore, limit cycles are less likely in our model compared to Webb et al. (2007b).

Our work also builds on previous studies of vertical transmission within lattice-based infectious disease models by including imperfect vertical transmission and both local and global transmission and reproduction. Silva et al. (2017) presents a lattice-based model with perfect vertical transmission and no recovery, and found that total infection is one possible outcome. We did not observe total infection in our model because imperfect vertical transmission and low levels of recovery are sufficient to maintain a susceptible host population even when horizontal transmission is high. As mentioned above, we found that there are horizontal transmission rates where vertical transmission is not a requirement for parasite invasion, in agreement with the 2-dimensional stochastic model of Schinazi (2000). Interestingly, their model did suggest that for a 1-dimensional lattice, if vertical transmission is below a critical threshold then regardless of the horizontal transmission rate the pathogen will not invade, suggesting that the precise details of the spatial structure may be crucial. A potential extension of our model would be to include spatial heterogeneity. For perfect vertical transmission, Su et al. (2019) found habitat loss to increase the importance of vertical transmission in disease prevalence. This is likely due to habitat loss decreasing the availability of susceptible hosts. If imperfect vertical transmission were included, then we expect that the availability of susceptible hosts would increase and that the pathogen could persist at higher levels of habitat loss.

**Fig. A1:**
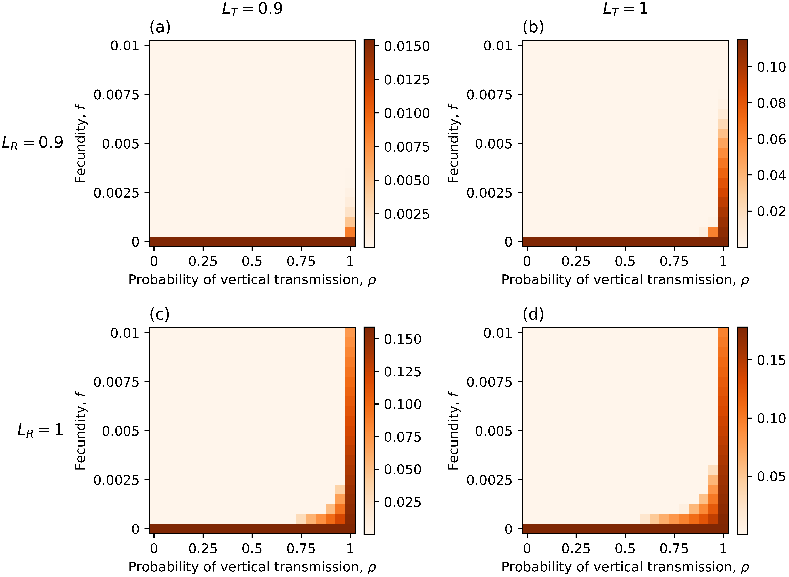
Heatmaps showing the amplitude of limit cycles for varying values of *L*_*T*_ and *L*_*R*_. The fixed parameters are *b* = 100, *d* = 1, *α* = 0.05, *γ* = 0.0005 and *β* = 100. The initial conditions are the same as Fig. 1

In summary, our findings show that vertical transmission can have a critical role in pathogen invasion and in the occurrence of limit cycles. Vertical transmission may also be the dominant cause of new cases when disease prevalence is high. Our results also suggest that evolving vertical transmission routes may be a favoured evolutionary strategy, which could be investigated further via an evolutionary spatial model.

## Acknowledgements

JW received funding from the Engineering and Physical Sciences Research Council (EPSRC).

## Appendix A

**Fecundity and Limit Cycles**

Fig. A1 shows that limit cycles only occur for very small values of *f* . When vertical transmission is perfect (*ρ* = 1), limit cycles occur for higher values of *f* compared to when it is imperfect. This suggests that infected hosts giving birth to susceptible hosts has as stabilising effect on limit cycles. This is consistent with results when there is no vertical transmission (Webb et al. 2007b).

## Appendix B

**Additional Equations for the Mutant Model**

### B.1 Mean-field Equations

The mean-field version of the spatial model described in section 3.2 is defined by the equations

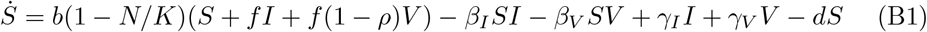

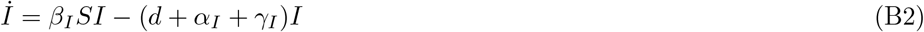

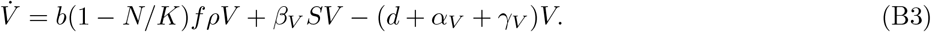

The parameters are as defined in Section 2, and, where applicable, the subscripts denote the strain corresponding to the parameter.

### B.2 Pair Equations

The corresponding pair equations for the system defined by Eq. 18-21 are

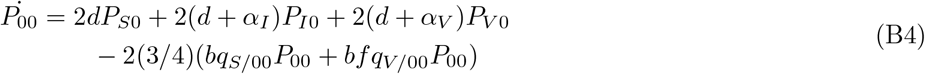

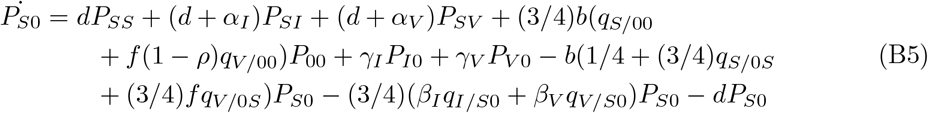

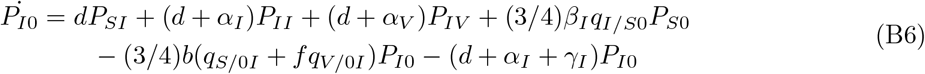

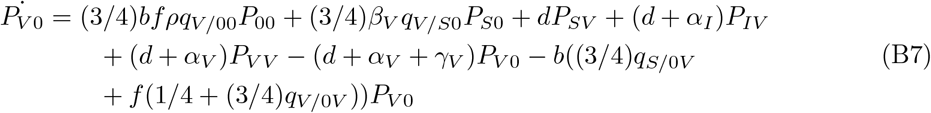

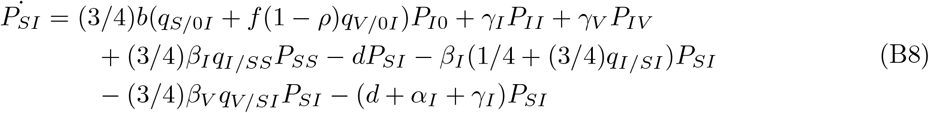

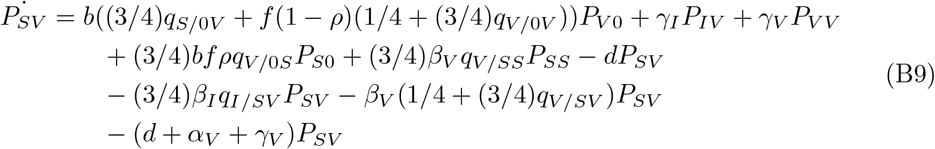

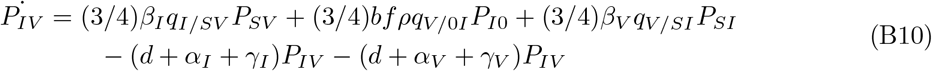

**Fig. C2:**
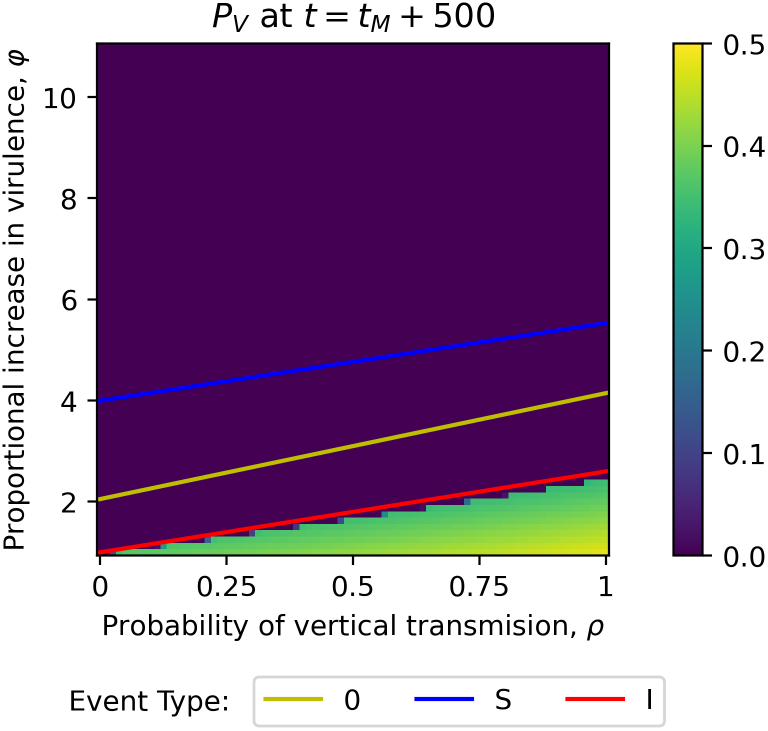
Heatmap showing *P*_*V*_ at *t* = *t*_*M*_ + 500 for the spatial model. The solid lines represent the boundary where 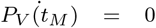. The fixed parameters are *b* = 10, *f* = 0.5, *d* = 1, *α*_*I*_ = 0.5, *α*_*V*_ = *φα*_*I*_, *γ*_*I*_ = *γ*_*V*_ = 1.5, *β*_*I*_ = *β*_*V*_ = 8, *t*_*M*_ = 500. The initial conditions are (*P*_0_, *P*_*S*_, *P*_*I*_, *P*_*V*_, *P*_00_, *P*_*S*0_, *P*_*I*0_, *P*_*V* 0_, *P*_*SI*_, *P*_*SV*_, *P*_*IV*_, *P*_*SS*_, *P*_*II*_, *P*_*V V*_) = (0.1, 0.8, 0.1, 0, 0.013, 0.078875, 0.008125, 0, 0.082375, 0, 0, 0.63875, 0.0095, 0) with 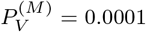

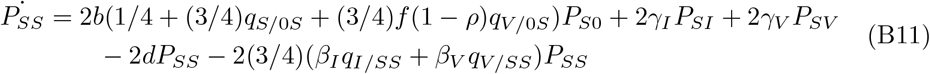

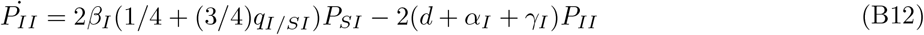

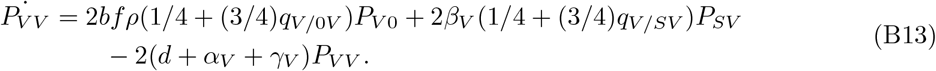

## Appendix C

**Virulence Trade-Off**

Here we consider the case when there is a trade-off between vertical transmission and virulence. We assume that the mutant strain has increased virulence, such that *α*_*V*_ = *φα*_*I*_ where *φ >* 1. As in the horizontal transmission trade-off case, for event type *S* and 0 the condition on 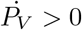 at *t*_*M*_ does not accurately predict the long term dynamics of the mutant. Whereas for event type 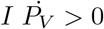 at *t*_*M*_ does predict the long term dynamics. Once again, we attribute this difference to the initial advantage in local environment of the mutant compared to the resident strain. As in the horizontal transmission trade-off case, the long-term dynamics are the same for all three event types.

## Statements and Declarations

### Author Contributions

Conceptualisation was by AB. Model derivation and analysis were performed by JW with supervision by B. The first draft of the manuscript was written by JW. Review and editing was done by both authors. All authors read and approved the final manuscript.

